# Genetic, epigenetic and metabolite variation in peripheral European Yew (*Taxus baccata* L.) populations at an unexplored part of the species natural distribution

**DOI:** 10.1101/2025.04.30.651400

**Authors:** Eleftheria Dalmaris, Evangelia Avramidou, Eirini Sarrou, Aliki Xanthopoulou, Salvatore Multari, Stefan Martens, Filippos Aravanopoulos

## Abstract

Taxanes form effective anticancer agents, which are found in the leaves and barks of the yew tree (*Taxus* L.). Taxol® (also known as paclitaxel), 10-diacetylbacatin III, 10-deacetyltaxol III, baccatin III and cephalommanine are anti-neoplastic taxanes used for cancer treatment. Due to the high demand of taxanes, it is of great pharmaceutical interest to investigate unexplored to date population diversity. In this context, three peripheral Greek *Taxus baccata* L. populations (Mt Cholomon, Mt Olympus and Mt Vourinos) were investigated to identify the extent and structure of their genetic (based on microsatellite markers), epigenetic (based on methylation sensitive amplified markers) and chemodiversity (based on liquid chromatography mass spectrometry) profiles. Results showed that the concentration of taxanes varied considerably in relation to population and harvest season. The main taxane in *T. baccata* needles was 10-deacetylbacatin III (DAB), with concentrations ranging from 267.8 (Mt Vourinos) to 517.6 (Mt Olympus) mg kg^-1^ dw. Besides metabolite variation, notable levels of genetic diversity and significant population differentiation were revealed. These results, in conjunction to the high levels of total methylation found in all populations, indicate their potential adaptability under climatic change. The findings of this study pave the way for prospective breeding and conservation strategies of these important *Taxus baccata* L. populations for artificial selection of highly producing taxane trees and protection of local germplasm.

## Introduction

Paclitaxel, the active ingredient of Taxol®, is an important bioactive compound found in the foliage and bark of several *Taxus* (yew) species [1]. The poisonous activity of the yew tree has been described since ancient times when Dioscorides, Pliny the Elder, Galen and Julius Cesar reported the “fortuitous accidents brought about by this poison [2]. The poet Shakespeare mentions the “magic power” of the yew tree in his play “Macbeth” (Macbeth, Act IV, Scene 1c). Paclitaxel is one of the three most used chemotherapeutic agents whose worldwide demand and price have increased exponentially [3, 4]. The paclitaxel biosynthetic pathway involves about 20 genes, and several primary metabolites [5], while a particular genomic region responsible for paclitaxel biosynthesis was recently found in the first complete *Taxus* genome published [6]. Originally paclitaxel (i.e., the active ingredient of Taxol®) was extracted from the bark of the Pacific yew tree (*Taxus brevifolia*), which proved to be unsustainable, as harvesting the bark ultimately kills the tree, while large amounts of bark are needed for small amounts of taxol [2, 7]. Nevertheless, although it is possible to chemically synthesize paclitaxel, the method is considered very expensive [8, 9]. Baccatin III and 10-deacetylbaccatin III have been identified as being important precursors of paclitaxel and are used for paclitaxel semi-synthesis [10, 11]. It is well documented that taxane concentration varies according to *Taxus* species or cultivar, location, season of sampling, the part of the tree that is used, and the age of the tree foliage [9, 12–18]. Furthermore, within and between populations, variation can also be high [19]. Considerable variation has been reported regarding the contents of paclitaxel, 10-deacetylbaccatin III and baccatin III, with ranges of 0-500 mg kg^1^, 0-4800 mg kg^-1^ and 0-500 mg kg^-1^ dried needles, respectively [20], which could provide a valuable germplasm for propagation and/or tissue culture experiments.

Paclitaxel is now mainly produced sustainably by semi-synthesis from precursor compounds, mainly from 10-deacetylbaccatin III found in the needles of European yew (*Taxus baccata*) [7, 21], a dioecious, wind-pollinated and animal-dispersed conifer, with scattered distribution in Europe and the Mediterranean [22]. To increase environmental sustainability and economic feasibility a few alternative taxol production methods are being investigated, including artificial cultivation in plantations [23–25]. Factors like geographical location, climate, and glaciation have influenced the genetic structure of *T. baccata* [26]. Results so far indicate a reduction of genetic diversity and an increase of genetic differentiation moving from northwest (central Europe, northern Iberian Peninsula) to southeast (Mediterranean Iberia and North Africa), in a study that covered the Western Mediterranean Basin range of *T. baccata* [27]. Furthermore, in recent years, studying DNA methylation in wild forest trees, became a prerequisite in order to understand plasticity, phenotypic variation, evolution, and adaptation to ongoing climatic changes [28–30]. Cytosine methylation is one of the most important epigenetic modifications in plants, playing a significant role in the regulation of gene activity [28], such as gene expression changes, secondary metabolite production and stress response [31]. In *T. baccata* it was shown that DNA methylation is involved in the regulation of paclitaxel biosynthesis [32]. Beyond variation in the genome and epigenome levels, chemodiversity especially regarding taxanes has also been seen. For instance, in three *Taxus* species, including *T. baccata*, distinct profiles were found by analyzing about 2,250 metabolites across various pathways, which emphasizes the genetic basis of chemodiversity and its impact on taxane biosynthesis [33]. Therefore, studying in a comprehensive manner the multifaceted genetic, epigenetic and chemodiversity of unexplored marginal and peripheral yew populations, is of high interest. Moreover, as European yew trees are highly toxic, they have been cleared by herders and farmers in many locations and have been decreased to small populations or become locally extinct [34]. As a result, the species has one of the highest declining rates in Europe. Therefore, the European Union has designated specially protected areas [Habitats Directive 92/43/EEC; 35]. In Greece, the protection of the species is limited, since only two populations have been identified as a priority habitat type according to the EC habitat’s directive (code 9580: “Mountainous coniferous woods with *Taxus baccata*”). Hence, a comprehensive diversity study is crucial not only for identifying future genetic resources for enhanced paclitaxel production, but as a mean to develop *in situ* and *ex situ* conservation strategies as well. Hence, the objectives of this study were: (a) to identify the chemodiversity within and between population of three peripheral Greek populations of *T. baccata,* focusing on five main taxanes, i.e., paclitaxel (PAC), 10-deacetylbaccatin III (DAB), baccatin III (BAC), 10-deacetyltaxol (DEAC), and cephalomannine (CEPH), (b) to study the seasonal variation of the three major taxanes i.e., paclitaxel, 10-daecetylbacatin III, and baccatin III, (c) to identify the extent and structure of within and between population genetic diversity, (d) to identify the levels of population epigenetic variation and the degree of DNA methylation, and (e) to provide background knowledge and guidelines for *in situ* and *ex situ* conservation strategies.

## Materials and Methods

### Experimental sites and collections

Leaf tissue was collected from 23-33 healthy *T. baccata* individuals at a minimum of 25 m tree to tree distance, from three natural populations: (1) Mt Cholomon (Chalkidiki), (2) Mt Olympus (Pieria) and (3) Mt Vourinos (Kozani; Fig 1 and S1 Table). Needle samples were collected at 2m height from north facing and well shaded branches. From each tree, tree height and diameter on breast height (DBH; S1 Fig) were recorded. Needle collection was conducted in three different seasons: (1) in early May, after the end of the flowering period (spring); (2) in late August, at the end of the growing season (summer) and (3) in early December, at the beginning of the winter period (winter).

**Fig 1.**
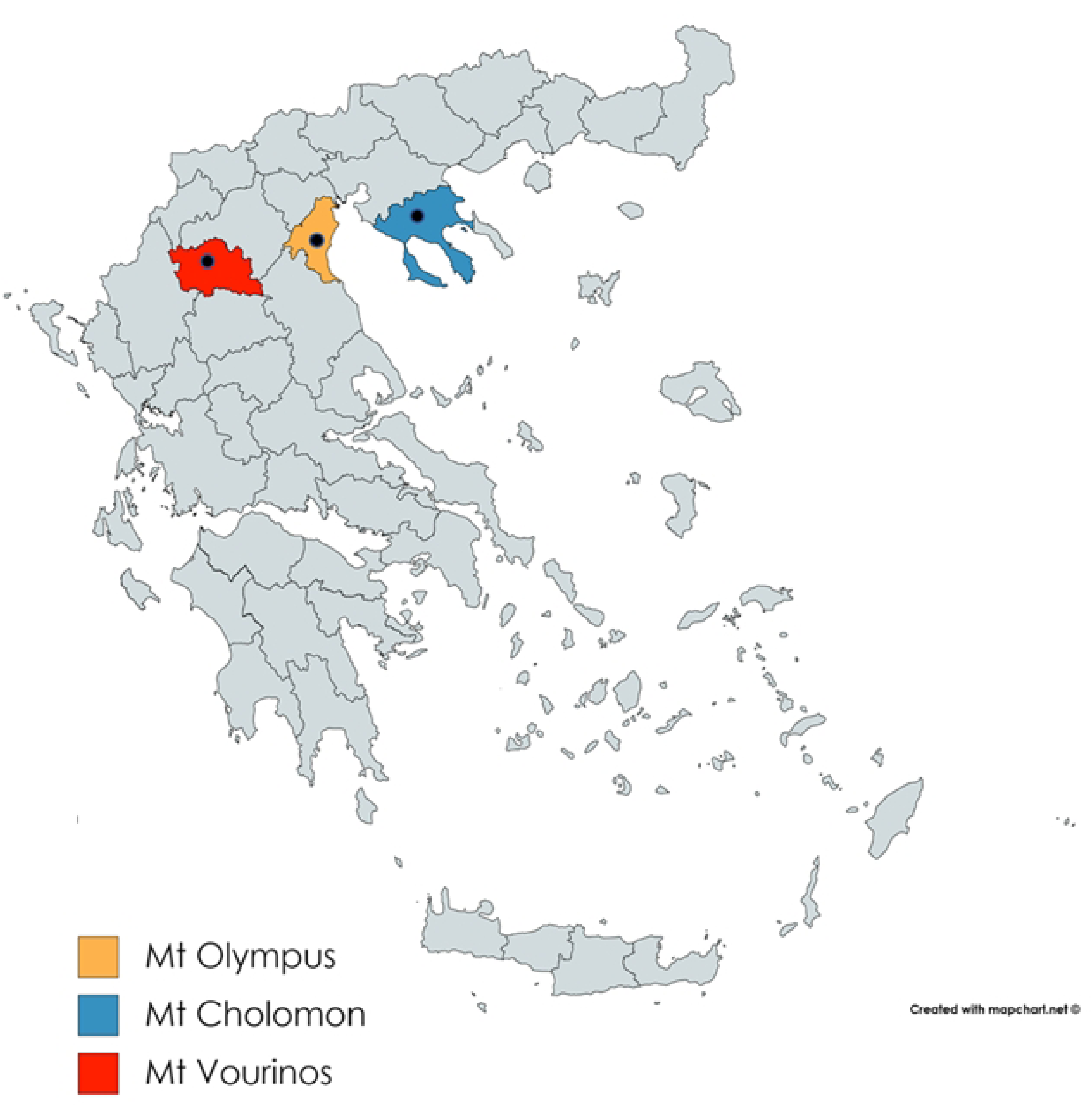
Locations of studied populations of *T. baccata* in Greece.

### Metabolomic analysis

#### Chemicals

Laboratory reagents were purchased from Sigma-Aldrich (Milan, Italy) and Fisher Scientific (Milan, Italy). Deionized water was purified *in loco* with the Arium® purification system (Sartorius AG, Goettingen, Germany). Authentic standards of taxanes, namely paclitaxel (PAC), 10-deacetylbaccatin III (DAB), baccatin III (BAC), 10-deacetyltaxol (DEAC), and cephalomannine (CEPH), were purchased from TransMIT (Gießen, Germany). Stock solutions of taxanes were prepared by dissolving the standards in methanol (LC-MS grade).

#### Taxanes extraction

The extraction of taxanes was performed according to the method of Mubeen et al. [36] with some modifications. The freshly collected needles were freeze-dried (Freeze-dryer Alpha 1-2 LD plus, Christ, Germany; at − 24 °C) and milled using a laboratory Mill IKA A11into homogenous powder. Samples (0.5 g) were mixed with 4 mL of 60% acetone, stirred on an orbital shaker for 20 mins, and centrifuged (5 min; 1800×*g*; 4°C) (procedure repeated thrice). The supernatants were pooled, and acetone was evaporated under reduced pressure at T ≤ 38 °C. The remaining aqueous phase was mixed with 4 mL of dichloromethane. The mixture was vortexed for about 1 min and centrifuged (2 min; 900×*g*; 4°C) (procedure repeated twice). The organic layers were collected and dried under reduced pressure at T ≤ 38 °C. The dry residue was dissolved in 2 mL of LC-MS methanol, filtered through PTFE 0.22 μm membranes into glass vials, and stored at -80 °C until further analysis. Results are expressed as mg kg^-1^ dw (dry weight).

#### Ultra Performance Liquid Chromatography Mass Spectrometry (UPLC-MS/MS)

The UPLC-MS analysis was adapted from Prokopowicz et al. [37] and performed on a UHPLC Dionex 3000 (Thermo Fisher Scientific Germany), equipped with a binary pump, an online vacuum degasser, and a column compartment. The separation of taxanes was carried out on a Kinetex C18 100Å column (2.1 mm × 100 mm, 2.6 μ), equipped with a column guard (Phenomenex, U.S.A.), kept at 35°C. Samples were injected using an autosampler (Dionex Thermo Fisher Scientific Germany) set at 4°C. The mobile phase was composed of water with 0.1 % v/v formic acid (A) and acetonitrile with 0.1 % v/v formic acid (B). Separation was carried out as follows: 80 % A (0–0.5 min), 80–50 % A (0.5–7 min), 50–28 % A (7–10 min), 20-0 % A (10-10.2 min), 0% A (10.2-12 min), 0-80% A (12.1-15 min). The flow rate was 0.3 mL min^−1^ and the injection volume was 5μL.

The MS/MS analysis was performed on an API 5500 triple-quadrupole mass spectrometer (Applied Biosystems/MDS Sciex, Toronto, Canada). The instrument was operated using an electrospray source in positive ion mode. The ESI parameters were as follows: the spray voltage was set at 5500 V for positive mode, the source temperature was set at 250 °C, the nebulizer gas (Gas 1) and heater gas (Gas 2) at 40 and 20 psi respectively. UHP nitrogen (99.999%) was used as both curtain and collision gas (CAD) at 20 and 9 psi respectively.) Analyst™ software version 1.6.1 (Applera Corporation, Norwalk, CT, USA) was used for instrument control and data acquisition. Compounds were identified by comparing the retention time and the spectral characteristic of the peaks with those of authentic standards. Multiple reactions monitoring (MRM) was used for quantification based on the peak area of the samples (Table 1).

**Table 1.**
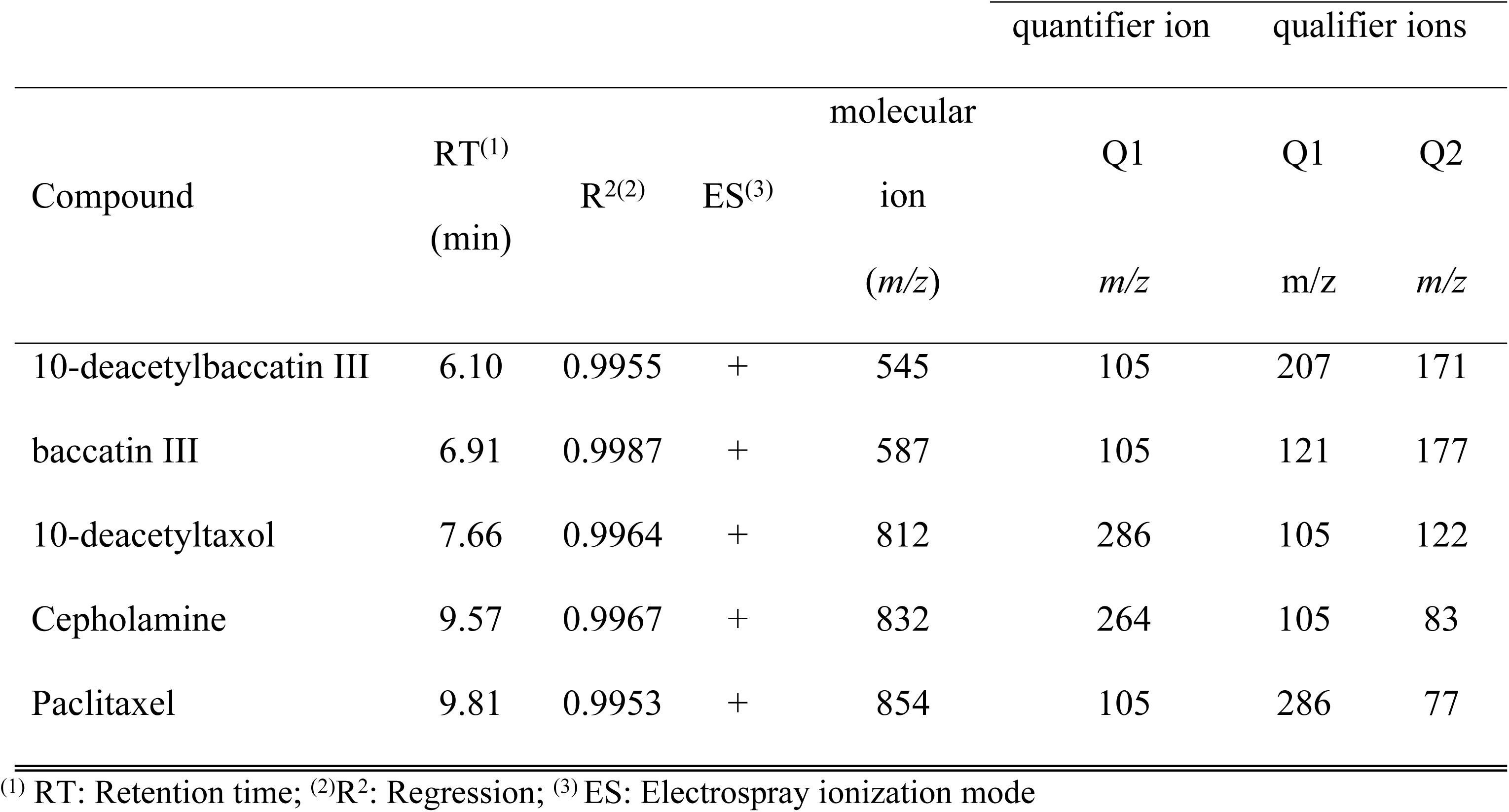
Multiple reactions monitoring (MRM) parameters of the five taxanes infused to the mass for optimization.

| Compound | RT<sup>(1)</sup><br>(min) | R<sup>2(2)</sup> | ES<sup>(3)</sup> | molecular<br>ion<br>(*m/z*) | quantifier ion | qualifier ions |  |
| --- | --- | --- | --- | --- | --- | --- | --- |
|  |  |  |  |  | Q1 | Q1 | Q2 |
|  |  |  |  |  | *m/z* | *m/z* | *m/z* |
| 10-deacetylbaccatin III | 6.10 | 0.9955 | + | 545 | 105 | 207 | 171 |
| baccatin III | 6.91 | 0.9987 | + | 587 | 105 | 121 | 177 |
| 10-deacetyltaxol | 7.66 | 0.9964 | + | 812 | 286 | 105 | 122 |
| Cepholamine | 9.57 | 0.9967 | + | 832 | 264 | 105 | 83 |
| Paclitaxel | 9.81 | 0.9953 | + | 854 | 105 | 286 | 77 |

<sup>(1)</sup> RT: Retention time; <sup>(2)</sup>R<sup>2</sup>: Regression; <sup>(3)</sup>ES: Electrospray ionization mode

### Microsatellite analysis

Twelve microsatellite loci were selected for genotyping: TAX23, TAX36, TAX60, TAX31, TAX09, TAX50, TAX26, TAX92, and TAX86 described by Dubreuil et al. [38], and: TB56, TB58, ΤΒ40 described by Mahmoodi et al. [39]. Polymerase chain reaction (PCR) conditions followed Dubreuil et al. [38] and Mahmoodi et al. [39], respectively. One μL of PCR product was added to 10 μL HiDi™ formamide (Applied Biosystems) and 0.15 μL 500 ROX Size Standard (Applied Biosystems), and capillary electrophoresis was run on an ABI3730 DNA Analyzer (Applied Biosystems). Results were analyzed using the GeneMapper Software v4.1 (Life Technologies), and allelic profiles were scored by automatic binning and visual checking.

### MSAP analysis

For the MSAP assay, double digests were initially performed using a combination of either *EcoRI/HpaII*, or *EcoRI/MspI* restriction enzymes. Digestion of 200-ng aliquots of genomic DNA was carried out in 20 μL containing 1Χ Cut smart buffer, 4 U *EcoRI* (New England, Biolabs) and 4 U of either *HpaII* or *Msp*I enzyme (New England, Biolabs) for 3 h at 37 °C. Two different adaptors designed to avoid reconstruction of restriction sites, one for the *EcoRI* sticky ends and one for the *HpaII/MspI* sticky ends, were ligated to the DNA after digestion, by adding to each final digestion 5 μL of a mix containing 5 pmol of EcoRI adapter, 50pmol of *HpaII/MspI* adapter, 1 mM ATP, 1Χ “One-for-All” Buffer, and 1 U of T4 DNA ligase (Invitrogen). The ligation mixture was incubated for 3 h at 25°C (S2 Table). Digested and ligated DNA fragments were diluted 5-fold and used as templates for the pre-selective amplification reaction. Pre-amplification reactions were performed using either *MspI/HpaII*-primers in a total volume of 2 μL containing 1X Kapa Taq Buffer, 0.2mM of each dNTP, 2.5 mM MgCl2, 30ng of each primer *EcoRI*+A, *MspI*/*HPaII*+A, 1U Taq DNA polymerase (Kapa Biosystems), and 5μL of diluted fragments (from the digestion and ligation reaction). Subsequently, pre-amplified fragments were diluted 10-fold and used as templates for the selective amplifications. For the selective amplification, only the *EcoRI* primers were labeled. Primer combinations employed and their sequences are shown in S2 Table. Selective PCR were performed in a 10μL total volume of 1X Kapa Taq Buffer, 2.5 mM MgCl2, 0.08mM of each dNTP, 5ng of labeled *EcoRI* primer, 30ng of HpaII/MspI primer, 1 U of Taq DNA polymerase (Kapa Biosystems), and 3μl of diluted pre-amplified DNA. Eight primer combinations were employed during the selective amplification stage. The whole experiment was repeated twice to only retain for further processing fully reproducible MSAP bands.

### Statistical Analysis

#### Metabolites

Analysis was processed using the R package [40]. A two-way-ANOVA analysis was performed. All data were checked for normality (by graphical data visualization and the Shapiro-Wilk test), and homogeneity of variances (by graphical data visualization and Levene’s test). Post hoc tests were conducted with Tukey’s HSD test method. A Biplot PCA analysis of metabolite profiling data derived from samples that were collected during the spring period performed using the XLSTAT statistical software (Version 2014.5.03; Addinsoft Inc., Brooklyn, NY, USA).

#### Microsatellites

Measures of intra-population and inter-population genetic parameters, number of observed (Na), and effective (Ne) alleles; observed (Ho) and expected (He) heterozygosity [41], were calculated using the GENEALEX 6.5 software [42, 43]. The inbreeding coefficient (F_IS_) for each population and each locus was obtained by computing a hierarchical AMOVA using the ARLEQUIN 3.5.2.2 software [44]. Statistical significance was determined by a non-parametric approach using 1000 permutations. As the assessment of allelic richness by the measure of allele frequencies needs to take into account the variation in population sizes, allelic richness (A) and the number of private alleles (Pa) were computed using the rarefaction method with HP-RARE software [45].

Analysis of molecular variance (AMOVA) to partition genetic variance was analysed in Arlequin 3.11. Principal coordinate analysis (PCoA) to analyze genetic structure by covariance standardized approach of pairwise [46] genetic distances was conducted in GenAlEx version 6.5. The relationships among populations were initially investigated by an unweighted pair group method using arithmetic means (UPGMA) dendrograms based on Nei’s [46] genetic distances. Nei’s genetic distance was calculated in GenAlEx v. 6.5. FigTree v1.3.1 was used to visualize and edit the tree. To delineate the genetic repartition of birch populations’ structure, a principal coordinate analysis (PCoA) was performed based on Nei’s unbiased genetic distance [41] using the GENEALEX 6.5 software.

To determine the genetic groups among populations, we used Bayesian clustering method in STRUCTURE V2.3.4. This analysis was run at 5 independent runs per K Value (K1–8) with a burn-in period of 100,000 iterations and 100,000 Markov chain Monte Carlo (MCMC). Structure Harvester was used to visualize the best K value based on delta K (ΔK). We used Mantel tests to determine the pattern of isolation by distance at 1000 permutations using GenAlEx version 6.537. The probability of identity (PID) was estimated in GenAlEx [42] to evaluate the discriminatory power of each locus. Deviations from Hardy-Weinberg equilibrium (HWD) were estimated using Genepop v4.2 [47].

#### MSAP

An AFLP Excel Macro [48] was used to convert allele size data from GeneMapper4.0 (Applied Biosystems, USA) into binary form and to indicate the presence “1” or absence “0” of fragments. Only reproducible fragments ranging from 150 to 500 bases were counted and further analyzed in order to reduce the impact of potential size homoplasy [49]. For MSAP analyses, comparison of the banding patterns of *EcoRI/HpaII* and *EcoRI/MspI* reactions, results in four conditions of a particular fragment: I: fragments present in both profiles (1/1), indicating an unmethylated state (n subepiloci); II: fragments present only in *EcoRI/MspI* profiles (0/1), indicating hemi or fully methylated CG-sites (h-subepiloci); III: fragments present only in *EcoRI/HpaII* profiles (1/0), indicating hemimethylated CHG-sites (m-subepiloci); and IV: absence of fragments in both profiles (0/0), representing an uninformative state caused either by different types of methylation, or due to restriction site polymorphism [50]. To separate unmethylated and methylated fragments and to test for the particular impact of the methylated conditions II and III, we used the ‘Mixed-Scoring 2’ approach [51].

Epigenetic diversity within populations was quantified using the R script MSAP_calc.r [51] as: (i) number of total and private bands (polymorphic subepiloci), (ii) percentage of polymorphic subepiloci (*P*epi) and (iii) mean Shannon’s information index (*I*epi). GenAlEx 6 [42] was employed to compute haploid gene diversity (*h*) within populations. GenAlEx was also used to conduct Analysis of Molecular Variance (AMOVA) - separately for each subepiloci class - in order to study the variation of CCGG methylation states (epiloci) among the five populations. A series of Principal Coordinate Analyses (PCoA) were employed to assess the ordination in multivariate space revealed by different types of epiloci.

## Results

### Dendrometric and climatic variables

The Mt Olympus population presented the tallest trees (7.9 – 31.4 m) and the largest DBH (35 – 115 cm) of all populations (S1 Fig), while the Mt Vourinos population had the shortest individuals (0.4 – 11 m) with the smallest DBH (0.2 – 20 cm) (S1 Fig). The latter population is grown in an area that presents the highest altitude and lowest annual precipitation of all populations (S1 Table). The other two populations grow at similar altitudes and precipitation levels.

#### Identification and seasonal variation of taxanes in *T. baccata* needles

A targeted LC-MS/MS method was adopted for the determination of the content of five taxanes, PAC, DAB, BAC, DEAC and CEPH. The phytochemical analysis conducted on an individual tree basis and the respective concentrations of the three populations are presented in S3 Table. Metabolite data from two populations (Mt Olympus and Mt Cholomon) have already been published [52]. Based on the samples that were collected during the spring period, DAB is the main taxane compound in the needles of the *T. baccata* trees, with concentrations varying from 267.8 (Mt Vourinos) to 517.6 (Mt Olympus) mg kg^-1^ dw. On the contrary, PAC, BAC, CEPH, and DEAC were detected in the extracts at lower amounts (12.9-24.4 mg kg^-1^ dw, 0.1-10.1 mg kg^-1^ dw, 11.8-16.2 mg kg^-1^ dw and 11.5-22.7 mg kg^-1^ dw, respectively). Significant differences were observed between populations (Table 2 and S4 Table). A biplot taxane metabolite analysis showed that the individuals from Mt Cholomon or Mt Vourinos populations are clustered closer, mainly covering two quartiles of the respective biplot, while the individual trees of Mt Olympus population revealed much higher dispersion (S2 Fig). In addition, none of the five taxanes determined in the samples, could characterize the populations as a discriminator compound. However, considering that: (i) DAB was the most predominant compound detected in the needle’s samples, (ii) PAC is the commercially valuable bioactive compound, and (iii) BAC serves as a precursor for the semi-synthesis of PAC, the investigation was carried out by focusing on these three taxanes out of the five originally analyzed.

**Table 2.** Statistical comparison of the *T. baccata* populations and Tukey HSD pairwise multiple comparison test.

| **(1) Significance after a two-way ANOVA** |  |  |  |
| --- | --- | --- | --- |
|  | **DAB** | **PAC** | **BAC** |
| **Season** | *** | ** | *** |
| **Pop** | *** | *** | *** |
| **Season \* Pop** | ** | * | *** |
| **(2) Tukey HSD pairwise multiple comparison test** |  |  |  |
| *i) For the Season effect* |  |  |  |
|  | **DAB** | **PAC** | **BAC** |
| **Spring** | A | A | A |
| **Summer** | B | A | B |
| **Winter** | B | B | C |
| *ii) For the Population* |  |  |  |
|  | **DAB** | **PAC** | **BAC** |
| **Mt Olympus** | A | A | B |
| **Mt Cholomon** | A | B | A |
| **Mt Vourinos** | B | C | B |

(1) Statistical comparison of the selected *T. baccata* populations for each of the three taxanes (DAB: 10 Deacetylbaccatin III, PAC: Paclitaxel, BAC: Baccatin III) after a two-way ANOVA with Populations (3 levels; Mt Olympus, Mt Cholomon and Mt Vourinos), Season (3 levels; Spring, Summer and Winter). Significant differences are represented with asterisks for $p \leq 0.000 - 0.001$ \*\*\*, $0.002 - 0.009$ \*\* and $0.01-0.05$ \*. (2) Populations and Season showed significant differences in all the measured compounds and therefore Tukey HSD pairwise multiple comparison tests were performed for the: (i) Season and (ii) Population. Sampling
period or population with a different uppercase letter are significantly different as determined by Tukey HSD pairwise multiple comparison test for $p \leq 0.05$ .

The concentration of DAB, PAC and BAC showed a significant seasonal variation (Fig 2, Table 2). The concentration of DAB differed significantly in spring samples in comparison to summer and winter ones, whereas PAC content in winter-collected samples varied significantly from spring and summer collection. Lastly, BAC concentration showed to be significantly different amongst all sampling periods (Fig 2 and Table 2). Besides seasonal variation, significant population variation was apparent as well. Samples from Mt Cholomon represented significantly higher concentrations for all three taxanes compared to samples from Mt Vourinos, whereas the BAC content was significantly higher compared only with Mt Olympus samples. The concentrations of DAB and PAC were significantly higher in samples from Mt Olympus compared to samples from Mt Vourinos (Fig 2 and Table 2).

**Fig 2.**
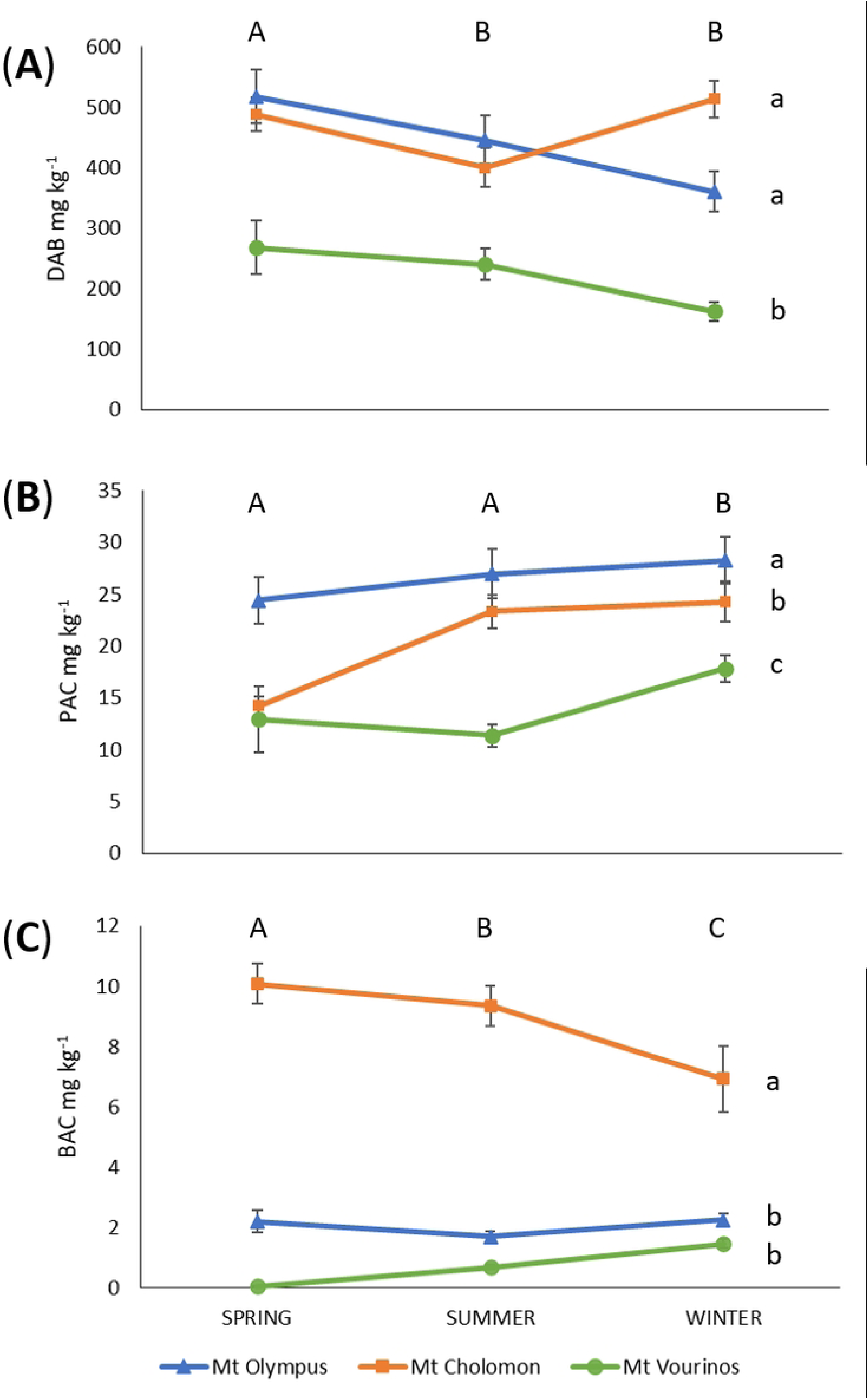
Seasonal variation of Paclitaxel, 10-Deacetylbaccatin III and Baccatin III in dry needles of *T. baccata* populations. (A) Paclitaxel (PAC), (B) 10-Deacetylbaccatin III (DAB), and (C) Baccatin III (BAC). Symbols represent the average of each population for the different compounds. Standard error bars represent variation within each of the populations. Sampling periods with a different uppercase (at top of each figure) and populations with a lowercase (at the side of each figure) letter are significantly different as determined by Tukey HSD pairwise multiple comparison test for *p* ≤ 0.05.

The concentration of all taxanes were categorized in three dissimilar patterns across populations and sampling periods (significant Season*Pop interactions, Table 2), but none of the taxanes followed a particular trend. The PAC content from Mt Olympus and Mt Cholomon needle samples and the BAC content from Mt Vourinos needle samples showed an increase in their concentration from spring to winter. On the other contrary, DAB concentration from Mt Olympus and Mt Vourinos samples, and BAC concentration from Mt Cholomon samples showed a decreasing trend from spring to winter. Lastly, the concentrations of PAC from Mt Vourinos, DAB from Mt Cholomon and BAC from Mt Olympus samples decreased from spring to summer, followed by an increase in the winter (Fig 2 and S3 Table).

Generally, the content of DAB in needles was always much greater than those of PAC and BAC, with the highest concentration being found in spring from the Mt Olympus populations (517.6 mg kg^-1^ dw). The highest PAC concentration was found also in the Mt Olympus population during the winter collection (28.3 mg kg^-1^ dw). Lastly, the maximum BAC concentration was found in Mt Cholomon samples during spring (S3 Table).

### Genetic variation of *T. baccata* populations

A total of 94 alleles were observed across the 12 SSR loci studied. The number of alleles per locus ranged from three (TB56) to 17 (TAX50), with a mean value of A=7.83 (S5 Table). The number of alleles per locus independent of sample size AR (the allelic richness) ranged from 4.167 (Mt Cholomon) to 6.083 (Mt Olympus) with a mean value of 5.000. Private alleles were also detected: private allelic richness (pAR) ranged from 0.583 to 1.500 with a mean 1.000, and on the average (pAR) amounted to about 15% of total AR. Observed heterozygosity ranged from 0.262 to 0.348 with a mean value of 0.299 for the three populations (S5 Table). The average expected heterozygosity (He) was highest in Mt Olympus (0.612) and lowest in Mt Cholomon (0.441; S5 Table). The probability of identity (P_ID_) was estimated at 1.6 x 10^-6^ for the Mt Cholomon samples and at 3.5 x 10^-10^ for the Mt Olympus genotypes whereas total P_ID_ was 1.5 x 10^-10^ (S5 Table). The inbreeding coefficient (Fis) ranged from 0.254 (TB40) to 1.000 (TAX36) with a mean value of 0.443 (S6 Table). At four loci (TAX23, TAX36, TAX09 and TAX50) the Fis value deviated significantly from zero (S6 Table).

Global genetic differentiation across all populations for SSR markers was estimated as F_ST_ and Gst (0.153 and 0.138, respectively). F_ST_ parameters varied among SSR loci, ranging from 0.004 (TB40) to 0.498 (TAX36; S6 Table). All pairwise Fst and Gst values were highly significant (*p* < 0.001; S6 Table). Gene differentiation coefficients (Gst, Fst) suggested significant population differentiation (moderately high Fst values, S7 Table). The AMOVA of the SSR data set produced congruent results (significant Φ statistics, Table 3). AMOVA also revealed a high percentage of variation due to population subdivision (76%). Furthermore, the inter-population AMOVA showed highly significant (*p* <0.001) genetic differences among populations. Thus, AMOVA (Φ_ST_ = 0.243; Table 3) also supports the results of F- and G-statistics (S6 Table). Furthermore, deviations from the Hardy-Weinberg equilibrium were found for all populations in many of the studied loci (S8 Table).

**Table 3.**
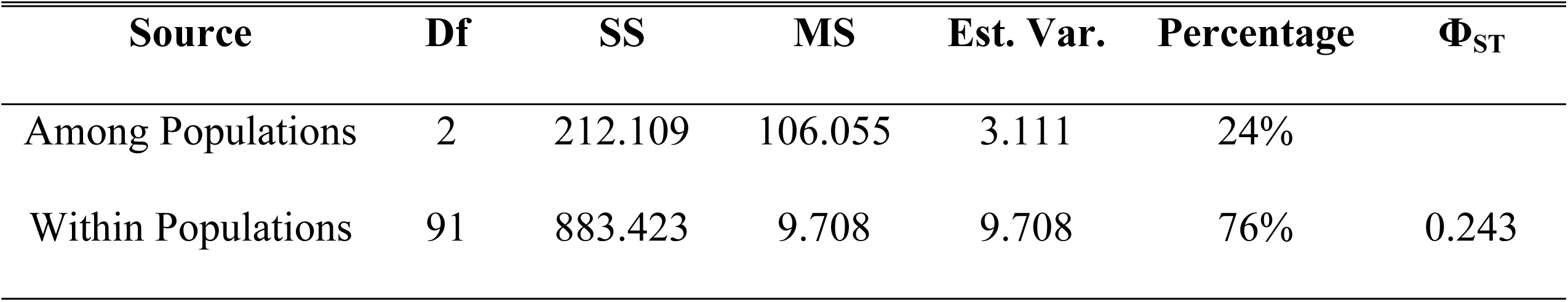
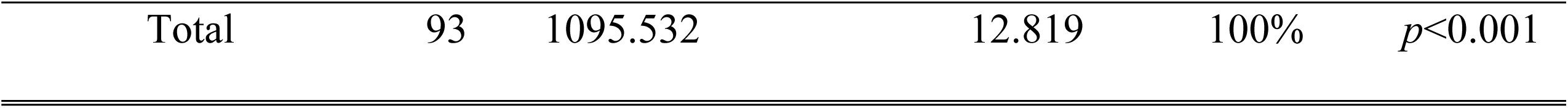
Analysis of molecular variance of the *T. baccata* populations based on SSR markers.

Relationships among populations were illustrated by a UPGMA Nei’s [46] genetic distance dendrogram, which grouped populations into two main clusters (S3 Fig). This finding was supported by PCoA (Fig 3) results which depicted two major clusters in the first two principal coordinates that accounted for 27.02% of the total variation (Fig 3). When a Mantel test was applied, there was a significant but weak correlation between geographic and genetic distances (*p* = 0.010, R^2^ = 0.140). These results were further corroborated by STRUCTURE analysis which was performed without prior information on the geographic origin of samples. The highest likelihood of the data was obtained for K=2. Low levels of admixture were observed in the three populations studied (Fig 4).

**Fig 3.**
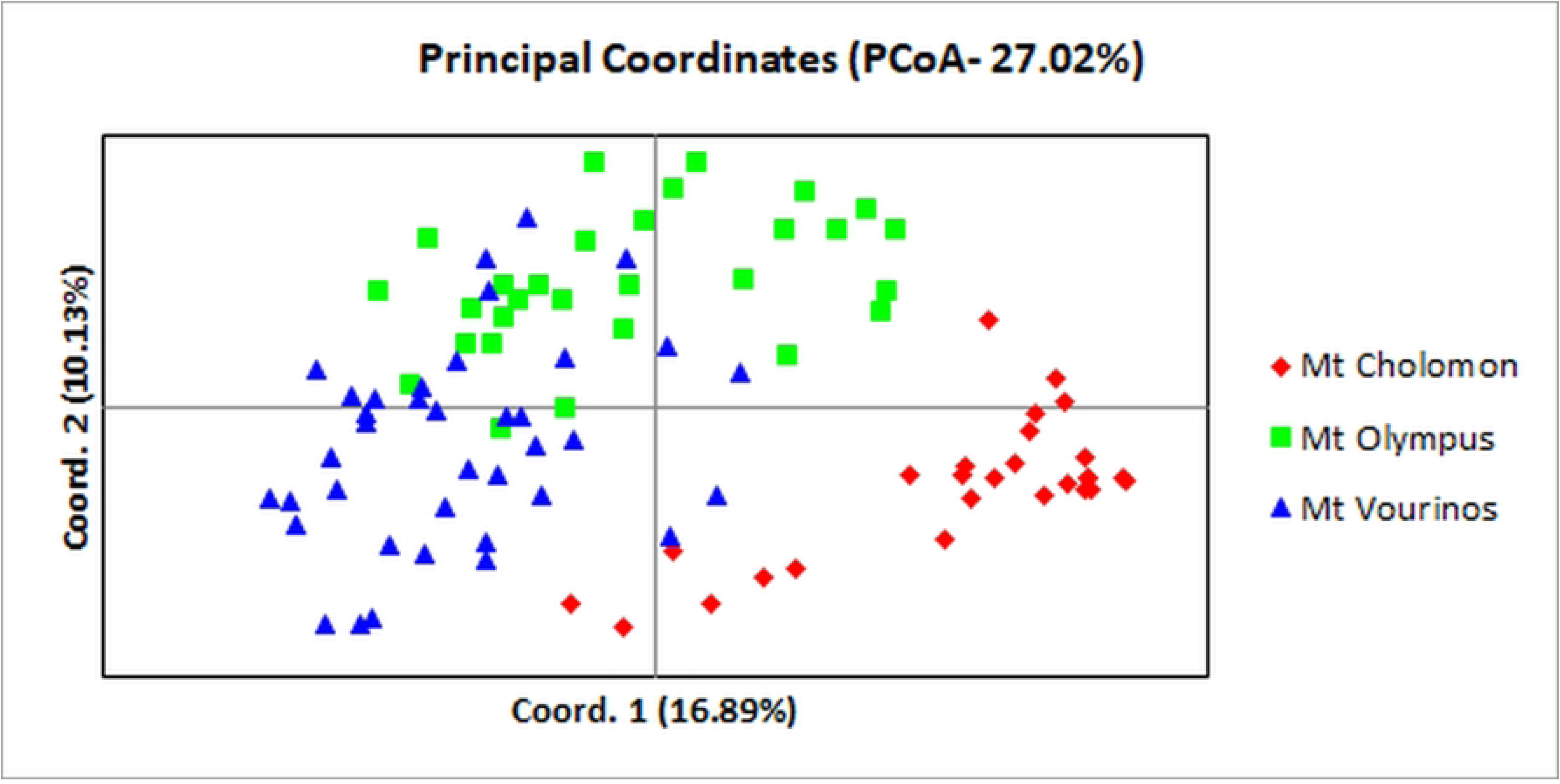
Principal coordinate analysis (PCoA) of 94 individuals in the *T. baccata* studied, based on 12 polymorphic SSR loci. A total of 27.02% of total variance accumulated on the first two components (axis 1 = 16.89%; axis 2 = 10.13%).

**Fig 4.**
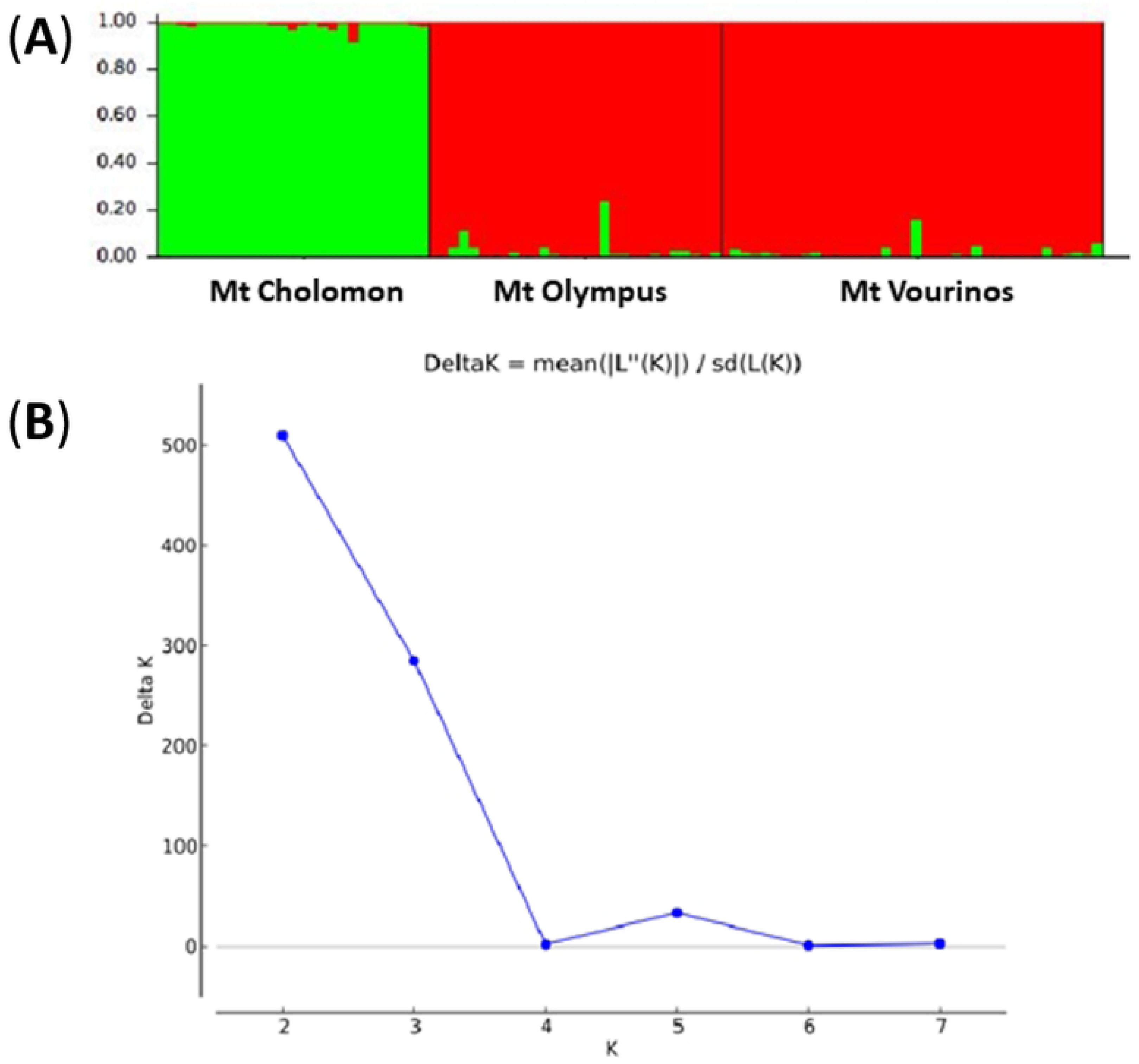
STRUCTURE analysis. (A) STRUCTURE analysis for SSR data with K= 2 clusters for the *T. baccata* populations studied. Barplots represent the average estimated membership probability (y axis) of an individual that belongs to a specific cluster. Each cluster is indicated by a different color. (B) Estimation of the number of populations for K ranging from 1 to 7 by calculating delta K values.

### Epigenetic variation of *T. baccata* populations

Epi-polymorphism varied among the populations studied (Table 4). The percentage of polymorphic epigenetic bands ranged from 46.16% in Mt Cholomon to 58.97% in Mt Olympus (mean percentage of polymorphic bands 53.21%). The number of private bands diversified from 462 (Mt Cholomon) to 865 (Mt Vourinos) (Table 4). The Shannon index (I_epi_), based on the frequency of methylation patterns within each marker type ranged from 0.188 (Mt Cholomon) to 0.223 (Mt Vourinos) (mean 0.209; Table 4). The Shannon index values were significantly different among the populations studied (Kruskal–Wallis χ^2^=24,588; *p*<0.001). The average index for methylation susceptible polymorphic loci was I_epi_=0.209).Lastly, MSAP mean haploid gene diversity (h) was 0.051 and ranged from 0.049 (Mt Cholomon) to 0.053 (Mt Olympus).

**Table 4.**
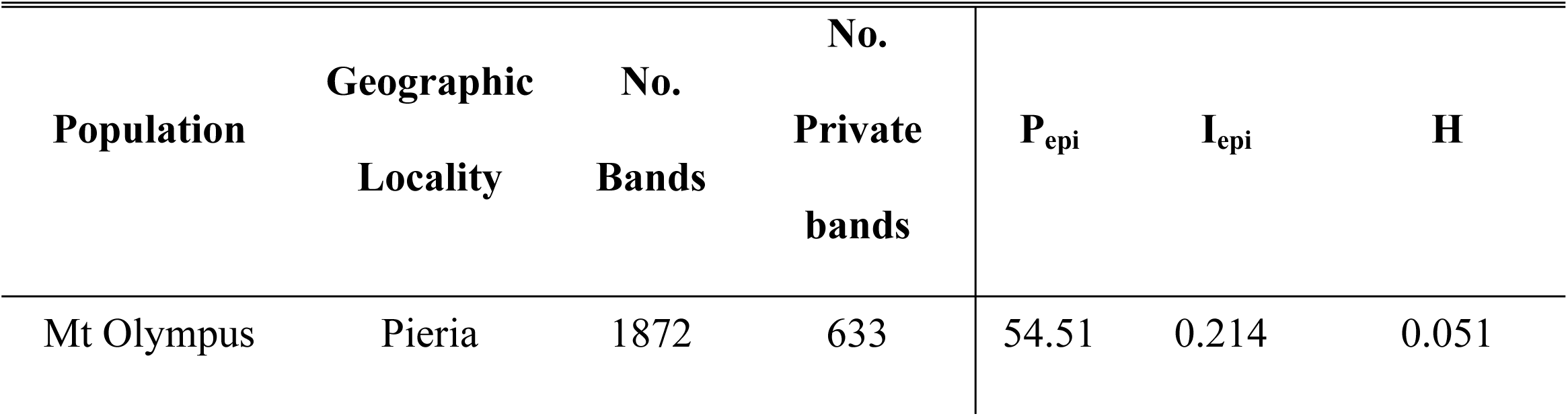
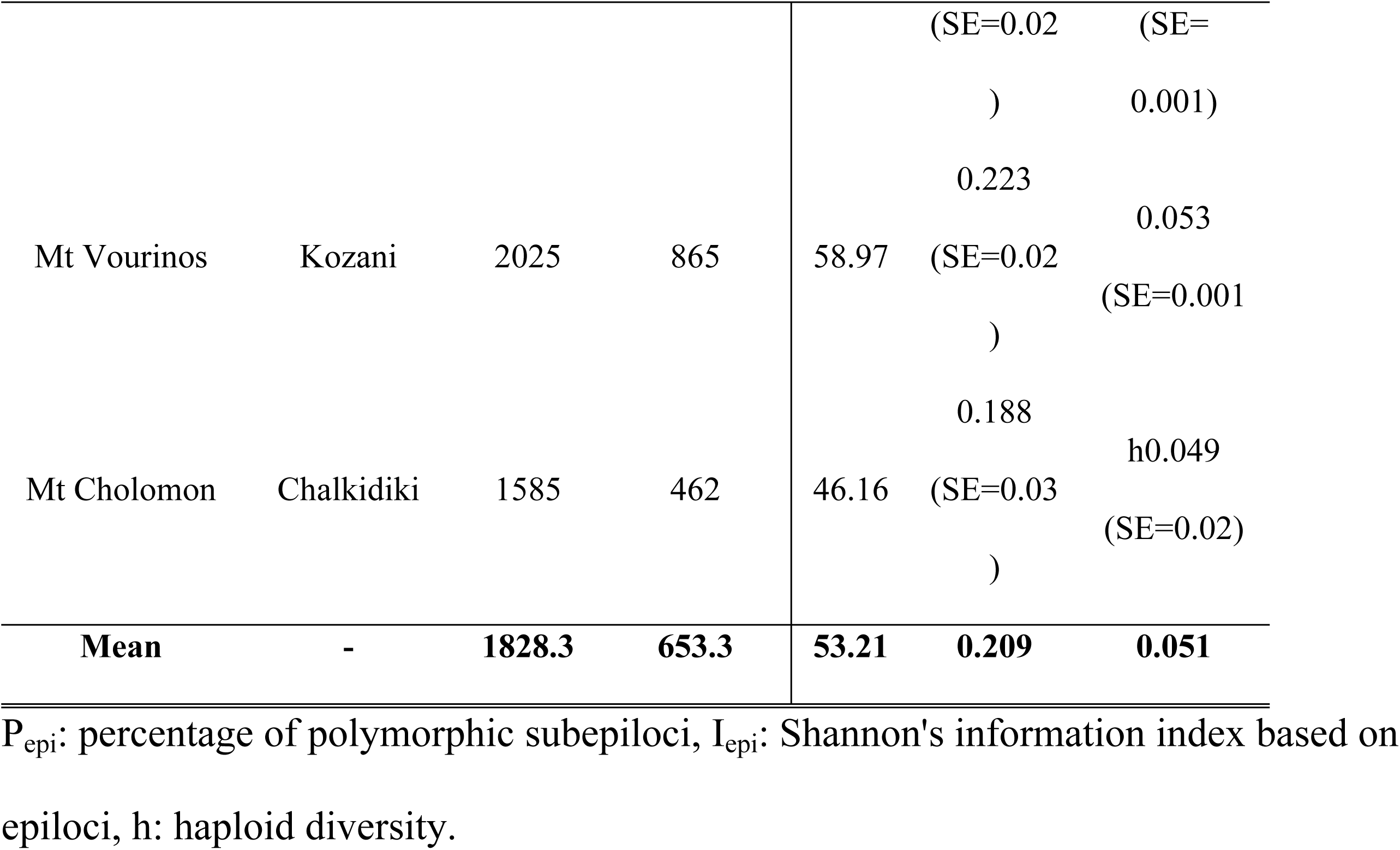
Total epigenetic diversity parameters (MSAP) of *T. baccata* populations.

AMOVA showed that approximately 86% of the total epigenetic variation was partitioned within populations (Table 5). Considering n-, m- and h-subepiloci separately, Φ_ST_ values were 0.168, 0.133, and 0.132, respectively (Table 5). The relative levels of full methylation (m-subepiloci), hemi-methylation (h-subepiloci) and non-methylation (n-subepiloci) presented an overall mean of 53.74%, 53.36 % and 51.19% respectively (Fig 5). The total level of 5’CCGG-methylation (sum of h- and m-subepiloci) ranged from 44% in Mt Cholomon to 62.33% in Mt Vourinos presenting (mean 53.55% across all populations). Population differences of 5’CCGG-methylation were not significant (Kruskal-Wallis x^2^=0.622 *p*=0.733).

**Fig 5.**
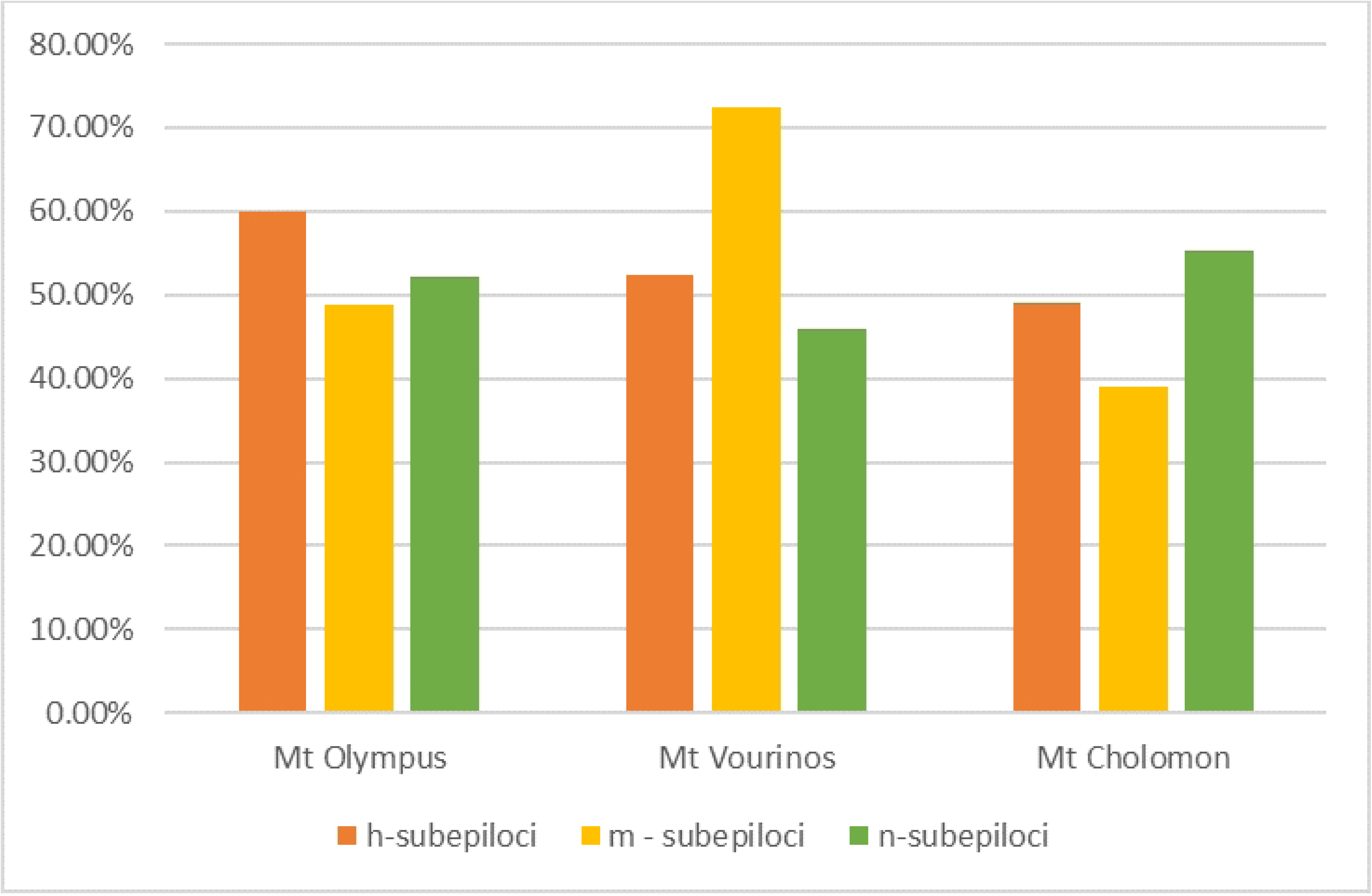
Percentage of polymorphic marker per population calculated from R script MSAP_calc.r for the *T. baccata* populations “h-subepiloci” “m-subepiloci” and “n-subepiloci”. Population differences of 5’CCGG-methylation were not significant (Kruskal-Wallis x2=0.622 *p*=0.733).

The PCoA of epigenetic distances revealed varying population differentiation in multivariate space (S4 Fig). The PCoA of m- and h-subepiloci failed to separate the populations and explained 21.92% and 20.5% of the total variation respectively. The same pattern was also depicted when all epi-alleles were combined and used for PCoA, where no clear differentiation of populations was evident and 20.32% of the total variation was explained.

Pairwise epigenetic F_ST_ values are being presented in Table S9. Epigenetic differentiation was moderate and ranged from 8.9% to 17.2%. When a Mantel test was applied, no significant relationship was found between the epigenetic distance and the geographical distance (R^2^=0.009, *p* < 0.020).

## Discussion

### Metabolomic analyses

The present study confirmed that the concentration of the investigated taxanes varied according to population (age, genetic pool, and site environmental conditions) and harvest season [14, 16, 17, 53]. The variance observed in the dendrometric and climatic variables of the populations under study (Table S1), contributed to this result. Taxol content depends on species, age and tissue type (e.g., needles, bark, stem) [9, 54] and therefore, caution needs to be taken when comparing results from the various studies which use different yew species protocols, class ages, tissue type, locations, sampling periods, etc. Overall, the concentrations of DAB, PAC, and BAC follow three different trends over the three collection periods. None of the taxanes showed a specific trend (Fig 2). Similar results indicating variation and different seasonal fluctuations throughout the year have been reported in several studies [14, 16, 17], whereas in other studies DAB, PAC, and BAC needle concentration reached their maximum at the same period e.g.: spring for *T. wallichiana* [15] or winter for *T. wallichiana* var. *mairei* [55]. Furthermore, DAB was measured to be the most abundant taxane of all and in all seasons in our study, which is in agreement with several other reports [16, 53, 55–57].

Overall, the main taxane in the needles of the yew trees studied was DAB and its highest concentration at the population level was found in Mt Olympus. DAB presented its maximum concentration in the needles collected at the end of the flowering period for Mt Olympus and Mt Vourinos populations, and at the beginning of winter for Mt Cholomon, whereas the maximum BAC needle concentration was found to be the opposite; in winter for Mt Olympus and Mt Vourinos and in spring for Mt Cholomon. Maximum concentration of DAB was found a month after the start of flush (June) in a seasonal study that used two cultivars (*T. x media* ‘Hicksii” and *T. x media* “Dark Green Spreader”; MI, USA) [58], whereas in the same study, BAC concentration of *T. x media* “Dark Green Spreader” had maximum concentration in the same period as DAB, but the maximum for the second cultivar was shown to be three months later, in September. In another seasonal study [17], August was identified to be the month with the highest DAB concentration, a month later than BAC concentration (Tehran Iran, *T. baccata*). in *T. wallichiana* needle samples from West Bengal India, the maximum concentration for both DAB and BAC was recorded during spring (March - April) [15]. Vance et al. [56] showed that both DAB and BAC needle concentration have the same optimum month for *T. brevifolia* (Oregon, USA), but that the maximum was identified to be in October. Lastly, *T. baccata* var. *fastigiate* DAB reached its maximum concentration in June (Ireland) [14] and in *T. canadensis* in August (New Brunswick, Canada) [16].

Needle samples obtained from the three *T. baccata* populations, accumulated higher PAC amounts in December at the beginning of winter when temperatures in Greece start to decrease, in contrast to the findings for the DAB concentration in the current study. Cameron & Smith [16] identified the highest PAC concentration in August-September, the same period as DAB (Canada, *T. canadensis*), Hook et al. [14] between February and April (*T. baccata* var*. fastigiate*), whereas Vance et al. [56] showed that *T. brevifolia* (Oregon, USA) PAC needle concentration expressed limited variation throughout an eight-month period (March to October), with a slight increase in late June. Lastly, the maximum PAC concentration was observed in June for *T. x media* ‘Hicksii” and *T. x media* “Dark Green Spreader” (MI, USA) [58].

Correlation analysis between the taxanes concentrations and climatic conditions (temperature, daylight etc.) has been employed to investigate whether these factors could somehow affect the *in-planta* biosynthesis and accumulation of the metabolites present in the needles [53, 55]. A positive and strong correlation was identified in *T. baccata* (Botanical Garden, Karaj, Iran) DAB needle and stem tissue concentrations with maximum, minimum and mean temperature, and light intensity of the day of sampling [53]. On the contrary, for BAC and PAC, no significant strong correlation with climatic conditions was identified [53]. Yang et al. [55], identified strong negativecorrelations between DAB needle concentration on one hand and monthly mean maximum temperature as well as length of daylight on the other in *T. wallichiana* var. *mairei* (Ningbo, China). The same study found similar correlation between PAC concentration and monthly mean maximum and minimum temperature [55}.

### Genetic analyses

At the population level, SSR markers indicated high levels of genetic variation in Mt Olympus and Mt Vourinos and relatively high levels in the population of Mt Cholomon. Notable amounts of genetic diversity were detected based on the observed (Ho=0.299) and expected (He=0.537) heterozygosity. The observed heterozygosity of the Greek populations was higher than that of populations from the Eastern Austrian Alps(Ho=0.178-0.272; [59]), lower in a population from Montseny Mountains in NE Spain (Ho=0.353; [60]), in populations from the northern part of the Czech Republic (H=0.42-0.57; [61]) and in western (Ho=0.508-0.622; [62]), and northern western (Ho=0.563-0.685; [63]) Polish populations, and lower that the average in a study that covered a large part of the natural range (195 populations, Ho=0.171-0.768, μ=0.474; [26]). The expected heterozygosity in populations from the Eastern Austrian Alps was found (similar to Ho) to be lower than our study (He=0.274; [59], whereas comparable values of He were discovered in NE Spanish (He=0.509; [60]) and western Polish (He=0.564; [62]) populations. Nevertheless, several other studies that analyzed populations of *T. baccata* from Spain, Britain, Poland and Czech Republic presented higher He values [27, 60, 61, 63–65], including a population from northern Poland that presented the highest He of all (He=0.870; [63]). Higher average He was also found in the study that covered a large part of the natural range (195 populations, He=0.406-0.855, μ=0.701; [26]). Furthermore, allelic richness (AR =5.000) of the Greek populations was within the range and higher than the average of the study that covered a large part of the distribution of *T. baccata* [195 populations (AR=2.243-5.295, μ=3.952; [26]) and higher than populations from Spain (AR=3.98, 4.23; [27, 65]) and Poland (AR=4.000; [62]).

Results indicate that there is less gene diversity as expected from HWE (Ho is lower than He) and some inbreeding may be involved, which is revealed by the positive value of Fis (0.443). Positive values of the inbreeding coefficient in *Taxus* species have been shown in several studies [60, 66–70]. The understory habitat of *Taxus* is considered as a factor promoting inbreeding, as it restricts pollen and seed dispersal thereby favoring mating between relatives [70, 71]. Also, the age structure of the studied Greek populations involves a relatively small number of old and large reproducing trees, a finding that may contribute to inbreeding.

F-, G-statistics and AMOVA are congruent and indicate moderately high genetic differentiation among populations (respective values of 0.153, 0.138 and 76% within population variation). Comparable results were shown in studies of *T. baccata* in the Western Mediterranean Basin (Fst= 0.155, 88% within population variation; [27]), in western Norway (Fst= 0.166; [68]), in northern Spain (Fst = 0.129, 88% within population variation; [65]) and in Poland (Fst = 0.155, 79.5% within population variation; [70]). Lower values of genetic differentiation were found in populations of *T. baccata* from Czech Republic (Fst = 0.068; [61]) and in British populations (Fst = 0.05, 63% within population variation; [64]).

### Epigenetic analyses

Studying methylation patterns in forest tree populations is becoming essential in order to access adaptation under climatic changes [28]. To the authors’ knowledge, this is the first study that involves the epigenetic MSAP method in natural populations of *Taxus* species. In one study MSAP have been employed in long term cell *Taxus media* cv. Hicksii cultures accompanied with HPLC analysis [32], and findings suggest that there was a higher level of DNA methylation in the low-paclitaxel yielding cell line for PAC biosynthesis after long-term culture. Wheeler et al. [19] studied five single trees from five natural populations of *T. brevifolia* with the TCL-HPLC method and suggested that taxol production is affected by epigenetic variation (e.g., time of collection, type of sampled material etc.).

In this study, the mean epigenetic diversity (Hepi) was 0.051, which was similar to that reported for Greek populations of the conifer *Pinus nigra* [Hepi=0.049; 72], but lower that Greek populations of the angiosperm *Prunus avium* populations [mean Hepi = 0.108; 73]. The total relative methylation in *Taxus* populations was 53.55% and was lower than *Pinus nigra* (68.02) but higher compared to *P. avium* Greek populations (49.67). In the above studies population differences are statistically significant, indicating the multifaceted level of methylation for natural populations. Total methylation was also assessed after vegetative propagation for *Pinus pinea* [74] and was higher compared to our results (64.36% vs. 53.55%). Such a difference could be due to several reasons, such as the different method used for scoring MSAP epiloci, the use of vegetatively propagated plants and genome wide methylation levels, which vary across individuals, plant organs and developmental stages.

### Comparative analysis of genetic and epigenetic diversity

In the present study, MSAP mean epigenetic diversity was h=0.051 and differed significantly (t=7.981, *p*=0.048) from SSR gene diversity (expected heterozygosity He=0.537). We have observed a negative non-significant correlation between genetic (SSR) and epigenetic (MSAP) diversity values regarding Hepi or h (r =−0.678, *p*=0.188) and Shannon’s epigenetic and genetic index (r=−0.693, *p*=0.163). Overall, genetic diversity (SSR) appears to be decoupled from epigenetic diversity (MSAP) in this study. These results should be treated with caution since the epigenetic markers uses are dominant, whereas the corresponding genetic markers are codominant. Albaladejo et al. [75] used MSAP and SSR markers in a greenhouse experiment for studying epigenetic and genetic correlation in *Pistacia lentiscus* between mother trees and offsprings. They also found that epigenetic variation was mostly decoupled from genetic variation.

## Conclusions

This study revealed a noteworthy diversity in within and between population taxane production. Based on the seasonal and between population chemodiversity of PAC, DAB, and BAC, and considering that DAB was found to be the prominent taxane, it can be concluded that the optimum time for harvesting is the spring, at the end of the flowering period in the Greek populations studied. Our results also provide evidence of adequate genetic variation and moderate population structuring, albeit with some inbreeding present in the peripheral small and fragmented populations that we studied. Nevertheless. total methylation levels were high, indicating a probable potential for future adaptation capacity under ongoing climatic change. Given the considerable between and within populations chemodiversity that was found, a selection of the high-end producing taxane trees is possible in the wild. Nevertheless given the fragile natural populations status, as well as environmental sustainability and economic feasibility concerns, the selected plus trees should not be directly harvested, but clonally propagated and established in artificial plantations in the frame of possible future marketable exploitation. Furthermore, considering the importance of *T. baccata* as the natural source of precursor compounds for the semi-synthetic paclitaxel production and the value of peripheral / marginal populations, it is important to expand the results of this study on the genomic, epigenomic and metabolomic profile of *T. baccata* in Greece.

The preservation of remaining *Taxus* populations in their natural environment and the conservation of their genetic resources is very important, especially given the rarity of yew trees in Greece, their fragmentation and their small population size (generally less than 50 individuals with low proportion of saplings and seedlings), but also their low representation under a statutory protection regime [34]. Conservation strategies should consider: (i) increasing the representation of protected areas that include *Taxus* woodlands, (ii) prohibiting yew logging, (iii) introducing genetic monitoring [76], (iv) regulating herbivory grazing and (v) if required, employ assisted gene flow [77]. *Ex-situ* conservation, using germplasm from different populations and potentially starting with the high-end producing taxane trees identified should certainly be considered in a comprehensive genetic conservation scheme.

## Acknowledgments

We would like to thank Anna-Maria Farsakoglou, Dr Ioannis Ganopoulos, Vasiliki-Maria Kotina, Theodoros Papadopoulos and Nikolaos Tourvas, for helping with field sampling.

## Supporting information

**S1 Fig. Height and DBH range of the *T. baccata* Mt Olympus, Mt Cholomon and Mt Vourinos populations.**

**S2 Fig. Biplot of LC-MS/MS analysis of taxane metabolite analysis for the *T. baccata* populations of Mt. Olympus, Mt Cholomon, and Mt Vourinos, based on the samples that were collected during the spring period.**

**S3 Fig. Genetic relationships of the populations of *T. baccata* based on SSR markers. Unweighted pair-group method using arithmetic average (UPGMA) unrooted tree illustrating genetic relationships among 94 individuals of *T. baccata* analyzed with 12 SSR loci. Samples included are indicated after population name.**

**S4 Fig. Principal Coordinate Analysis of epigenetic data. PCoA are partitioned into three methylation types: u alleles, m alleles, h alleles and all epigenetic alleles.**

**S1 Table. Altitude (m), annual rainfall (mm) and coordinates of the *T. baccata* populations employed in this study.**

**S2 Table. Primers and adapters used in the f-MSAP analysis of the *T. baccata* populations**

**S3 Table. Descriptive statistics of the concentration of 10 Deacetylbaccatin III (DAB), Paclitaxel (PAC), Baccatin III (BAC), Cephalomannine (CEPH) and10-deacetylTaxol (10DEAC) of dry shaded needles in the T. baccata populations studied.**

**S4 Table. Statistical comparison of the *T. baccata* populations and Tukey HSD pairwise multiple comparison test for cephalomannine (CEPH) and 10-deacetylTaxol (10DEAC). S5 Τable. Genetic diversity of the *T. baccata* populations of this study.**

**S6 Τable. F- and G-statistics for 12 SSR loci analyzed in the *T. baccata* populations studied.**

**S7 Table. Genetic Pairwise Fst Values in the *T. baccata* populations studied.**

**S8 Table. Summary of Chi-Square Tests for Hardy-Weinberg Equilibrium of the *T. baccata* populations studied.**

**S9 Table. Epigenetic Pairwise F_ST_ Values in the *T. baccata* populations studied.**

